# The Red Rice Bran Extract (RRBE) Mitigates Photoaging by Targeting Mitochondrial Oxidative Stress and Regulating Thermal Damage Responses

**DOI:** 10.64898/2026.01.08.698301

**Authors:** Ting-Syuan Lin, Yigang Chen, Xin Li, Xiang Ji, Shenghan Huang, Jiaying Lyu, Liping Li, Huali Zuo, Shangfu Li, Jing Li, Hsi-Yuan Huang, Hongxin Zhuo, Xintao Zhang, Yong-Fei Wang, Shangqian Ning, Zhihua Zhang, Ye Gu, Tao Zhang, Yang-Chi-Dung Lin, Hsien-Da Huang

## Abstract

The photoprotective efficacy of natural skin-active complexes is well recognized, yet their application is often hindered by the challenge of deciphering their complex, multi-component, and multi-target mechanisms. To bridge the gap between established phenotypes and molecular mechanisms, we developed an AI-driven integrated platform that combines phytochemical profiling, network pharmacology, and deep learning-based target prediction with rigorous biophysical validation. We applied this platform to investigate RRBE, a bioactive complex refined from red rice bran extract. *In vivo* clinical studies confirmed that RRBE significantly accelerates the resolution of UV-induced erythema, while cellular and 3D tissue models demonstrated robust suppression of oxidative stress and DNA damage responses. To decode its material basis, the platform deconstructed RRBE into 10 distinct chemical modules. Leveraging our SCOPE-DTI deep learning model for global target prediction, we identified flavonoids (Module 1) and phenolic acids (Module 10) as the primary bioactive drivers relevant to photoprotection. These computational predictions were structurally supported by molecular docking and definitively validated in a physiological environment via Cellular Thermal Shift Assay-Mass Spectrometry (CETSA-MS). Mechanistically, RRBE functions through a synergistic polypharmacology: (1) Module 1 components, represented by procyanidin B2, target NDUFA7 to stabilize mitochondrial function and mitigate ROS; (2) Module 10 components, exemplified by caffeic acid, ferulic acid, and p-coumaric acid, engage FKBP11 and HSP90AA1 to regulate protein homeostasis and stress responses. This work not only deciphers the polypharmacological basis of RRBE’s photoprotective action but also validates a scalable “AI-guided, cell-validated” discovery pipeline, offering a rational paradigm for uncovering the protective benefits of complex natural extracts.

## 1 Introduction

Skin aging is a complex biological phenomenon characterized by a progressive decline in tissue function and regenerative capacity, driven by both intrinsic and extrinsic factors [1]. Chronic exposure to solar ultraviolet (UV) radiation is the most significant extrinsic contributor, leading to photoaging, a pathological process that accelerates skin deterioration far beyond chronological aging [2].

The UV spectrum consists of UVA (320–400 nm) and UVB (290–320 nm), which inflict damage through distinct mechanisms. UVA penetrates deeply into the dermis, where it primarily generates oxidative stress that leads to collagen degradation, while the more energetic UVB radiation is absorbed by the superficial epidermis, causing direct DNA mutations and inflammation. This cumulative damage manifests clinically as deep wrinkles (rhytides), rough texture, mottled hyperpigmentation, and a profound loss of skin elasticity. Beyond its aesthetic impact, photoaging compromises the skin’s critical barrier function and is mechanistically linked to photocarcinogenesis. UV radiation is classified as a “complete carcinogen,” [3], acting as both a tumor initiator and promoter, which significantly increases the risk of precancerous lesions and skin cancers, including melanoma and non-melanoma types such as basal cell and squamous cell carcinomas.

At the heart of UV-induced photoaging is the excessive production of reactive oxygen species (ROS), which, at elevated levels, overwhelm the skin’s antioxidant defenses, causing oxidative stress [4]. ROS molecules damage lipids, proteins, and DNA, triggering destructive signaling cascades. This oxidative stress activates pro-inflammatory pathways, including NF-*κ*B and MAPK signaling [5], leading to the upregulation of matrix metalloproteinases (MMPs) and chronic inflammation [6].

Crucially, solar exposure involves not only oxidative insult but also thermal and proteotoxic stress. The energy absorbed from UV and infrared radiation generates heat, disrupting cellular protein homeostasis (proteostasis) [7]. This thermal stress, combined with ROS-induced oxidation, leads to the accumulation of misfolded proteins in the endoplasmic reticulum (ER), triggering ER stress and the Unfolded Protein Response (UPR) [8]. To maintain survival, skin cells rely on molecular chaperones, such as Heat Shock Proteins (HSPs) and immunophilins (e.g., FKBPs), to refold damaged proteins and resolve inflammation [9]. Therefore, effective anti-photoaging strategies must not only scavenge ROS but also bolster the skin’s intrinsic capacity to manage thermal and protein folding stress.

Mitochondria are both a major source of ROS and a vulnerable target of this multifaceted stress. Mitochondrial dysfunction, particularly impaired electron transport chain (ETC) function, results in a “ROS burst” that further exacerbates proteotoxicity and inflammation [10]. Importantly, oxidative stress-induced mitochondrial dysfunction activates NF-*κ*B, creating a detrimental feedback loop that compromises cellular homeostasis [11]. Thus, targeting mitochondrial integrity and the associated stress response networks is pivotal for preventing photoaging.

Natural products, derived from plants, fungi, and microorganisms, have long been a source of therapeutic agents. Many natural complexes exhibit polypharmacology, modulating multiple biological targets simultaneously—such as antioxidant enzymes and stress response chaperones—making them ideal for treating complex conditions like photoaging [12]. However, natural product research has faced significant challenges due to the multi-component nature of these compounds, complicating the elucidation of their mechanisms of action [13]. The advent of AI and machine learning has revolutionized this field, enabling efficient data analysis and prediction of synergistic interactions [14]. This computational approach, known as network pharmacology, has reinvigorated natural product research and opened new pathways for drug discovery [15].

Among these natural sources, extracts derived from rice bran, the nutrient-rich outer layer of the rice grain, have been utilized for centuries in traditional skincare [16]. Rice bran is a rich source of bioactive compounds, including polyphenols, peptides, and vitamins. Specifically, pigmented varieties such as red rice contain high concentrations of flavonoids and phenolic acids, which exhibit significant anti-aging effects [17]. While studies have shown these compounds can inhibit MMPs and protect against UV damage [18], their specific molecular targets—especially regarding mitochondrial and thermal stress regulation—remain largely unexplored.

To bridge the gap between the macroscopic photoprotective effects of natural products and their molecular drivers, we developed a “digital-to-cellular” framework that couples *In Silico* target prediction with cell-based validation applied to RRBE, a red rice bran extract (Figure 1). Integrating phytochemical profiling with our previously developed deep learning model, SCOPE-DTI, we predicted the direct targets of RRBE’s key components and validated these interactions using Cellular Thermal Shift Assay-Mass Spectrometry (CETSA-MS). This study is the first to deconvolute RRBE’s efficacy into specific compound-target axes, identifying NDUFA7 as a mitochondrial guardian and FKBP11/HSP90AA1 as modulators of proteostasis. These insights provide a robust molecular basis for RRBE’s dual action against oxidative and thermal stress, establishing a new paradigm for evidence-based research on natural products.

**Figure 1:**
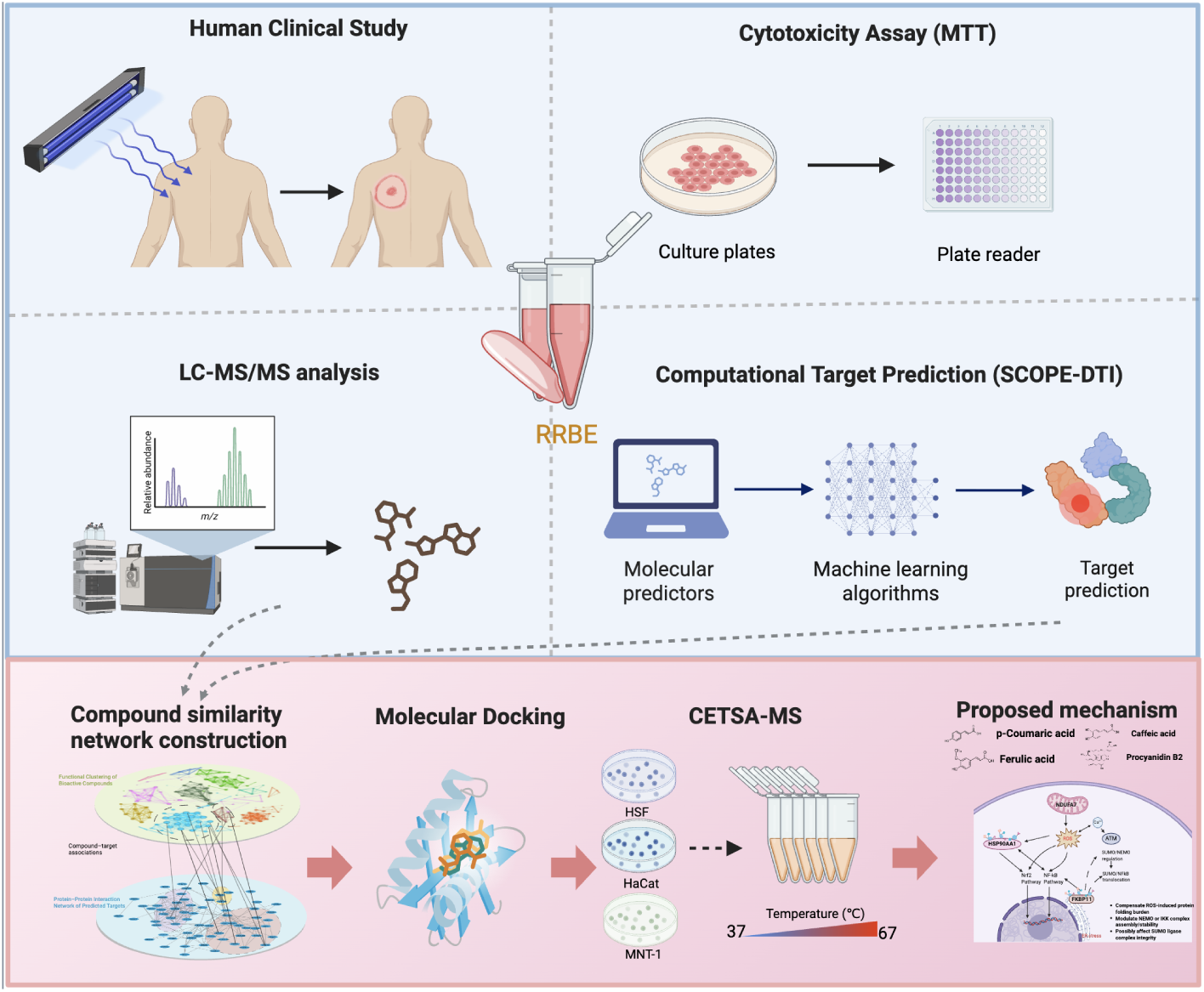
The workflow of this study. The integrated platform bridges the phenotypic effect of RRBE with molecular mechanisms by utilizing LC-MS/MS profiling, AI-driven target prediction (SCOPE-DTI), and experimental validation (CETSA-MS).

## 2 Materials and Methods

### 2.1 Materials and Reagents

The standardized red rice bran extract (RRBE, Lot No. EPSP240924) was provided by Better Way (Shanghai) Cosmetics Co., Ltd. Reference standards (purity *>* 98%) including caffeic acid, ferulic acid, p-coumaric acid, and procyanidin B2 were purchased from Aladdin (Shanghai, China). Recombinant human proteins (NDUFA7, FKBP11, and HSP90AA1) were obtained from Abnova. Primary antibodies against Cyclobutane Pyrimidine Dimers (CPDs) (Cosmo Bio), 8-OHdG (Abcam), and NF-*κ*B (p65) (Abcam) were used for immunodetection.

For *in vivo* and 3D tissue evaluations, specific dermatological formulations were prepared. The investigational product (RRBE Lotion) was formulated by incorporating 1% (w/w) of the RRBE extract. Based on the raw material’s solid content of 0.68%, the final concentration of active RRBE solids in the formulation was 68 ppm. A matching Negative Control (Vehicle) was prepared using the identical base formulation without the active extract. The Positive Control consisted of a 7% Ascorbic Acid (Vitamin C) formulation in a silicone-stabilized system to ensure stability. The complete ingredient lists for all three formulations are detailed in Supplementary Table S1.

### 2.2 Human Clinical Study (Photoprotection Assay)

The *in vivo* photoprotective efficacy study was conducted on 10 healthy adult volunteers (*n* = 10), aged 18–60, after standard screening. Prior to participation, all subjects provided written informed consent, and the study adhered to the Declaration of Helsinki guidelines. Approval was granted by the Ethics Committee of Better Way (Shanghai) Cosmetics Co., Ltd. (Approval No. BWG/2025-03). The Minimal Erythema Dose (MED) was determined for each subject one day prior to the main test. On the test day, four distinct 49 cm*^2^* (7×7 cm) sites on the subjects’ backs were randomly assigned to one of four groups: Control (UV), Positive Control (7% Vitamin C + UV), Negative Control (vehicle + UV), and RRBE (68 ppm RRBE solids + UV). Test samples were applied at a standardized density of 2.0 ± 0.1 mg/cm*^2^*. Twenty minutes post-application, to allow for sample absorption and film formation, the designated sites were irradiated with 2.5× MED using a UV lamp. Skin erythema was instrumentally quantified using a Mexameter^®^ to record the Erythema Index (E.I.) at baseline and at 72 hours (72 h) post-irradiation.

### 2.3 Cell Culture

Human epidermal keratinocytes (HaCaT) and human dermal fibroblasts (HSF) were obtained from Newgainbio Co., Ltd., while the human melanoma cell line (MNT-1) was purchased from the American Type Culture Collection (ATCC, Manassas, VA, USA). HaCaT cells were cultured in MEM (Gibco, Billings, MT, USA, No. 31095029), whereas HSF and MNT-1 cells were maintained in DMEM (Gibco, No. C11995500BT). The media for HaCaT and HSF were supplemented with 10% FBS (Gibco, No. 26170043). In contrast, the medium for MNT-1 was supplemented with 20% FBS (Gibco, No. 26170043) and 1% NEAA (Gibco, No. 11140-050). All culture media contained 1% Penicillin-Streptomycin. All cells were maintained in a humidified incubator at 37^◦^C with 5% CO*_2_*.

### 2.4 Evaluation of Photoprotection in 3D Reconstructed Human Epidermal Models

#### 2.4.1 Model Culture, Irradiation, and Treatment

The 3D reconstructed human epidermal models were transferred to 6-well plates containing 0.9 mL of pre-warmed EpiGrowth culture medium per well. The models were divided into four groups: Blank Control (BC), Negative Control (NC), Positive Control (PC), and RRBE-treated group. Except for the BC group, all models were exposed to UVB irradiation at a dose of 600 mJ/cm*^2^* for approximately 2 min 29 s.

Immediately following irradiation, treatment was administered. For the PC group, the positive control agents (Vitamin E, Dexamethasone, or WY-14643, depending on the assay) were added directly into the culture medium underneath the insert. For the RRBE group, 25 *µ*L of a working solution containing 68 ppm RRBE solids (in EpiGrowth medium) was applied topically and distributed evenly onto the surface of the epidermal model. The models were then incubated at 37^◦^C in a 5% CO*_2_* humidified atmosphere for 24 h. After incubation, the model surfaces were rinsed with sterile PBS, and residual liquid was gently removed. The tissues were then fixed in 4% paraformaldehyde (PFA) for 24 h prior to histological processing.

#### 2.4.2 Immunohistochemical (IHC) Analysis of CPDs

To assess direct DNA damage, paraffin-embedded tissue sections were prepared. The sections were deparaffinized, rehydrated, and subjected to antigen retrieval. They were then incubated with a primary antibody against Cyclobutane Pyrimidine Dimers (CPDs) overnight at 4^◦^C, followed by incubation with a horseradish peroxidase (HRP)-conjugated secondary antibody. DAB (3,3’-diaminobenzidine) was used as the chromogen to visualize CPDs (brown staining). Images were captured under a light microscope, and the Mean Integrated Optical Density (Mean IOD) of the nuclear staining was analyzed to quantify DNA damage.

#### 2.4.3 Immunofluorescence (IF) Analysis of 8-OHdG

Oxidative DNA damage was evaluated by detecting 8-hydroxy-2’-deoxyguanosine (8-OHdG). Fixed tissue sections underwent antigen retrieval and blocking, followed by incubation with anti-8-OHdG primary antibody. A fluorophore-conjugated secondary antibody was used for detection. Nuclei were counterstained with DAPI. Fluorescence images were acquired using a fluorescence microscope. The levels of 8-OHdG were quantified by measuring the Relative Fluorescence Intensity, normalized to the Blank Control.

#### 2.4.4 Immunofluorescence (IF) Analysis of NF-*κ*B

To evaluate the inflammatory response, the expression and localization of NF-*κ*B p65 were determined using immunofluorescence. Tissue sections were incubated with anti-NF-*κ*B p65 primary antibody, followed by the appropriate fluorescent secondary antibody and DAPI counterstaining. The intensity of NF-*κ*B fluorescence was captured and analyzed. The relative expression levels were quantified as Relative Fluorescence Intensity compared to the control groups.

#### 2.4.5 Histological Quantification of Sunburn Cells (SBCs)

To assess acute phototoxicity and apoptosis, tissue sections were stained with Hematoxylin and Eosin (H&E). Sunburn cells (SBCs) were identified morphologically by their characteristic pyknotic nuclei and eosinophilic cytoplasm. The number of SBCs was counted in randomly selected fields of view under a light microscope. The degree of tissue damage was expressed as the mean number of SBCs per field.

### 2.5 Measurement of UVA-Induced Intracellular ROS

To assess protection against UVA-induced oxidative stress, HaCaT keratinocytes were seeded in 96-well plates and pre-treated with RRBE for 24 h. Cells were then washed and incubated with 10 *µ*M DCFH-DA probe (2’,7’-dichlorofluorescein diacetate) for 20 min at 37^◦^C. Following probe loading, cells were exposed to UVA irradiation to induce ROS generation. Intracellular ROS levels were quantified by measuring the fluorescence intensity of DCF (Excitation: 480 nm; Emission: 525 nm) using a fluorescence microplate reader.

### 2.6 Gene Expression Analysis via Quantitative Real-Time PCR (RT-qPCR)

#### 2.6.1 Cell Treatment and Experimental Design

HaCaT cells were revived and seeded into 6-well plates. For p53 analysis, cells were treated when they reached 30%–50% confluence. The cells were divided into Blank Control (BC), Negative Control (NC), Positive Control (PC, Vitamin E), and RRBE-treated groups. After 24 h of treatment, all groups except the BC were exposed to UVB irradiation at a dose of 300 mJ/cm*^2^* (irradiation time: 1 min 14 s), followed by an additional 24 h incubation at 37^◦^C with 5% CO*_2_*. For HSF1 and HSP70 analysis, cells were treated when they reached 40%–60% confluence.

The groups (BC, PC, and RRBE) were treated with their respective agents (with Vitamin E serving as the PC) for 24 h at 37^◦^C with 5% CO*_2_* without UVB irradiation, to assess the direct modulatory effects of RRBE on stress response gene expression.

#### 2.6.2 RNA Extraction and Quantitative Analysis

Following the respective incubation periods, cells were washed twice with PBS. Total RNA was extracted using RNAiso Plus (1 mL/well) according to the manufacturer’s protocol. The RNA was reverse-transcribed into cDNA. Gene expression levels of *p53*, *HSF1*, and *HSP70* were quantified using fluorescence quantitative PCR (qPCR). The specific primer sequences used in this study are listed in Table 1. Gene expression was normalized using *β-actin* as the internal reference gene.

**Table 1:**
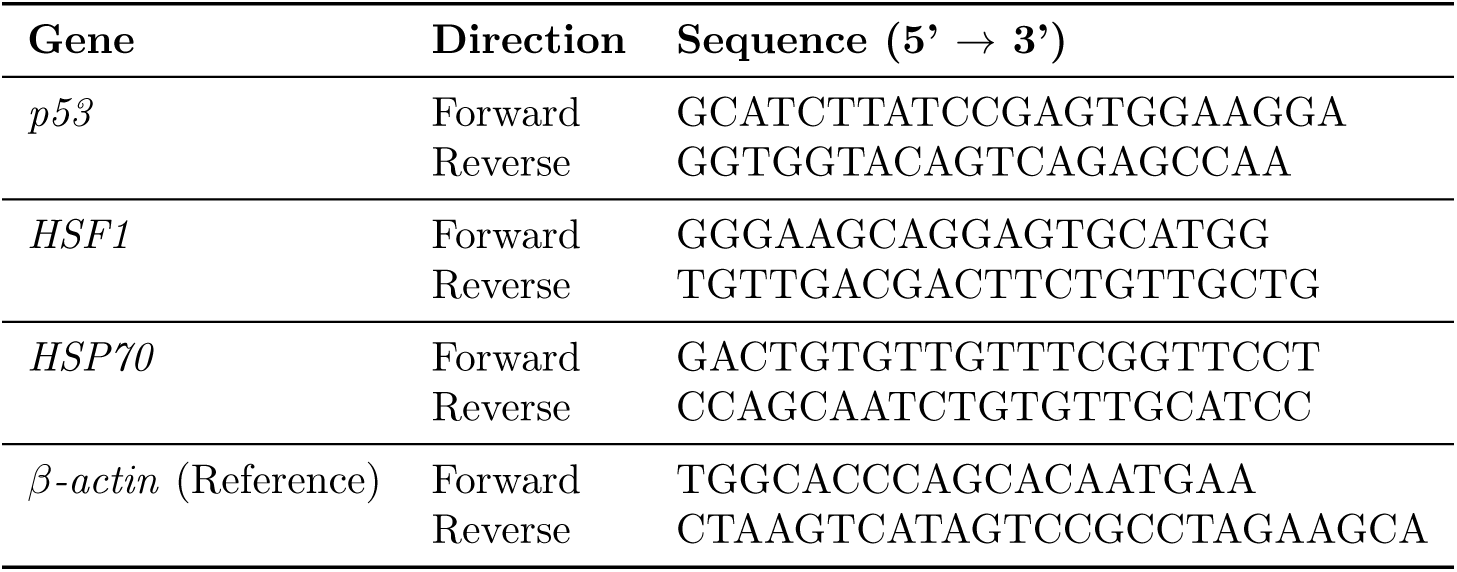
Primer sequences used for RT-qPCR.

### 2.7 Cytotoxicity Assay (MTT)

HaCaT cells were seeded in 96-well plates to 50-60% confluency and treated in triplicate (*n* = 3) with RRBE (0.078125% to 10%, v/v) or controls (Solvent, 10% DMSO) for 24 h. MTT solution (0.5 mg/mL) was added for 4 h, then replaced with 150 *µ*L of DMSO. Absorbance was read at 490 nm.

### 2.8 LC-MS/MS Analysis

**UHPLC:** Samples were analyzed using an Ultra-High Performance Liquid Chromatography (UHPLC) (Shimadzu Nexera UHPLC LC-30A) system coupled with a mass spectrometer (TripleTOF 6600+, Sciex). Separation was performed on a C18 column (2 *µ*m C18, 2.1 × 100 mm). Mobile phase A comprised 99.9% water and 0.01% formic acid, while mobile phase B was 100% acetonitrile. Gradient elution was conducted with the following program: 0-2 min, from 0% to 5% B; 2-22 min, 99% B; 22-26 min, 99% B; 26-30 min, from 99% to 5% B. Flow rate: 0.3 mL/min; column temperature: 30^◦^C; injection volume: 20 *µ*L.

**MS:** Metabolites were identified by MS using a mass spectrometer (TripleTOF 6600+, Sciex) after being separated by UHPLC. The ion scan range (m/z) was 100–2000 Da, and the analysis was conducted employing SWATH acquisition in both positive and negative ion modes. The parameters were set as follows: curtain gas (CUR) set to 35 psi; gas source 1 (GS1) at 50 psi; gas source 2 (GS2) at 50 psi; ion source temperature (TEM) at 550^◦^C; MS1 accumulation time at 250 ms; MS2 accumulation time at 100 ms. For the positive mode, the ion spray voltage (ISVF) is set at 5500 V, while for the negative mode, the ISVF is set at -4500 V.

#### Data Processing and Relative Quantification

Raw SWATH data were processed using MS-DIAL version 5.4. The following settings were applied: MS1 tolerance = 0.01 Da, MS2 tolerance = 0.025 Da, retention time window = 0–25 min, MS1 mass range = 100–2000 Da, MS2 mass range = 50–2000 Da, minimum peak height = 500 amplitude, MS/MS abundance cut-off = 5 amplitude. Metabolite identification was based on matching with the MS-DIAL internal database, using an MS1 tolerance of 0.01 Da and an MS2 tolerance of 0.05 Da. Alignment was performed using pooled QC samples as reference (retention time tolerance = 0.05 min, MS1 tolerance = 0.015 Da). The relative abundance of each metabolite was represented by its peak area.

### 2.9 Compound Similarity (CS) Network Construction

The compound similarity searches were conducted using RDKit based on molecular fingerprints. A compound similarity (CS) network was constructed, where nodes represent individual compounds and edges represent the relationships between them. Compounds with a Tanimoto coefficient (Tc) ≥ 0.5 were considered connected in the network. To identify and separate the clusters within the CS network, the Louvain algorithm in Gephi was applied with a resolution of 1.0. Additionally, independent structural analyses were performed on the nodes (compounds) within each cluster using chemical data obtained from PubChem.

### 2.10 Computational Target Prediction (SCOPE-DTI)

The molecular targets for the 210 compounds identified by LC-MS/MS were predicted using SCOPE-DTI, a deep learning framework for predicting drug-target interactions (DTIs). The SMILES structures for each compound were used as input to the model. The resulting ranked list of predicted protein targets for each compound was then utilized for the subsequent module clustering and pathway enrichment analyses. The detailed architecture, training, and validation of the SCOPE-DTI model are described in the original publication [19].

### 2.11 Pathway Importance Analysis

The calculation of functional dimension importance for each module follows a weighted approach that considers both protein importance and the prevalence of functions. The core calculation process involves the following steps:

Each protein’s importance is calculated using min-max normalization of its count value:

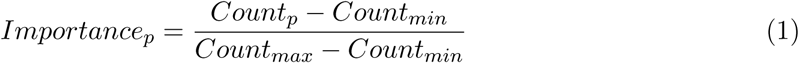

Where *Count_p_* is the count value of protein *p*, and *Count_max_* and *Count_min_* are the minimum and maximum count values within that module.

For each functional dimension *d*, two factors are calculated:

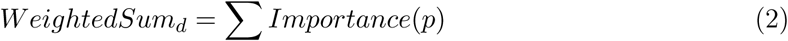

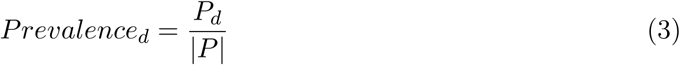

Where *P_d_* is the number of proteins with function *d*, and | *P* | is the total number of proteins in the module.

The final importance score for each dimension combines both factors:

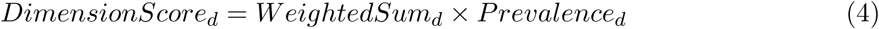

### 2.12 Molecular Docking

Protein–ligand docking was conducted using AutoDock Vina (version 1.1.2) [20]. Protein structures were preprocessed by removing crystallographic water molecules and adding polar hydrogens. Ligands were converted to docking-ready formats using MGLTools and Open Babel [21]. Binding pockets were identified with P2Rank [22].

### 2.13 Cellular Thermal Shift Assay-Mass Spectrometry (CETSA-MS)

#### 2.13.1 Preparation of Cell Lysate for Target Identification

The collected cell pellets were suspended in PBS containing a mixture of 1% protease inhibitors. The cell lysate was extracted by performing three rapid freeze-thaw cycles in liquid nitrogen. The mixture was centrifuged at 20,000 g for 20 minutes at 4^◦^C. The supernatant concentration was determined by the BCA method and diluted to 1 mg/mL with lysis buffer.

#### 2.13.2 Target Identification

The sample was prepared following the method developed by Lyu et al. [23] with minor modifications. Briefly, cell lysate was divided into two parts and treated with compounds (10 *µ*mol/L) or DMSO for 10 min. Sera-Mag carboxylate-modified magnetic particles were added (mass ratio 5:1). Samples were incubated at 52^◦^C for 3 min, then cooled. Magnetic beads were separated, washed, and subjected to alkylation and trypsin digestion overnight.

## LC-MS/MS Analysis

Peptide samples were subjected to SWATH-MS based label-free quantitative proteomics using Ekspert nanoLC 400 coupled with AB Sciex TripleTOF 6600 plus. Separation was performed on a C18 column using a 90-minute gradient.

## Data Processing

Raw data was processed using DIA-NN 1.9.1 [24] against a spectral library of >10,000 human proteins [25]. Differential expression analyses were performed using Limma [26].

## 3 Results

### 3.1 Clinical Efficacy and Mechanistic Investigation of RRBE in Mitigating UV-Induced Skin Damage

#### 3.1.1 RRBE Significantly Accelerates the Resolution of UV-Induced Erythema in Human Subjects

To objectively quantify the *in vivo* photoprotective efficacy of RRBE, a clinical study (*n* = 10) was conducted by irradiating skin sites with 2.5× the Minimal Erythema Dose (MED). Skin erythema was instrumentally monitored using a Mexameter, measuring the absolute Erythema Index (E.I.) at baseline and 72 hours post-irradiation to assess the resolution of inflammation.

Following UVB exposure, the Negative Control group exhibited a sustained inflammatory response, maintaining a high mean E.I. level of ∼267.9 at 72 hours (Figure 2). This persistent erythema indicates that the vehicle alone failed to effectively mitigate the UV-induced vascular damage within the observation period.

**Figure 2:**
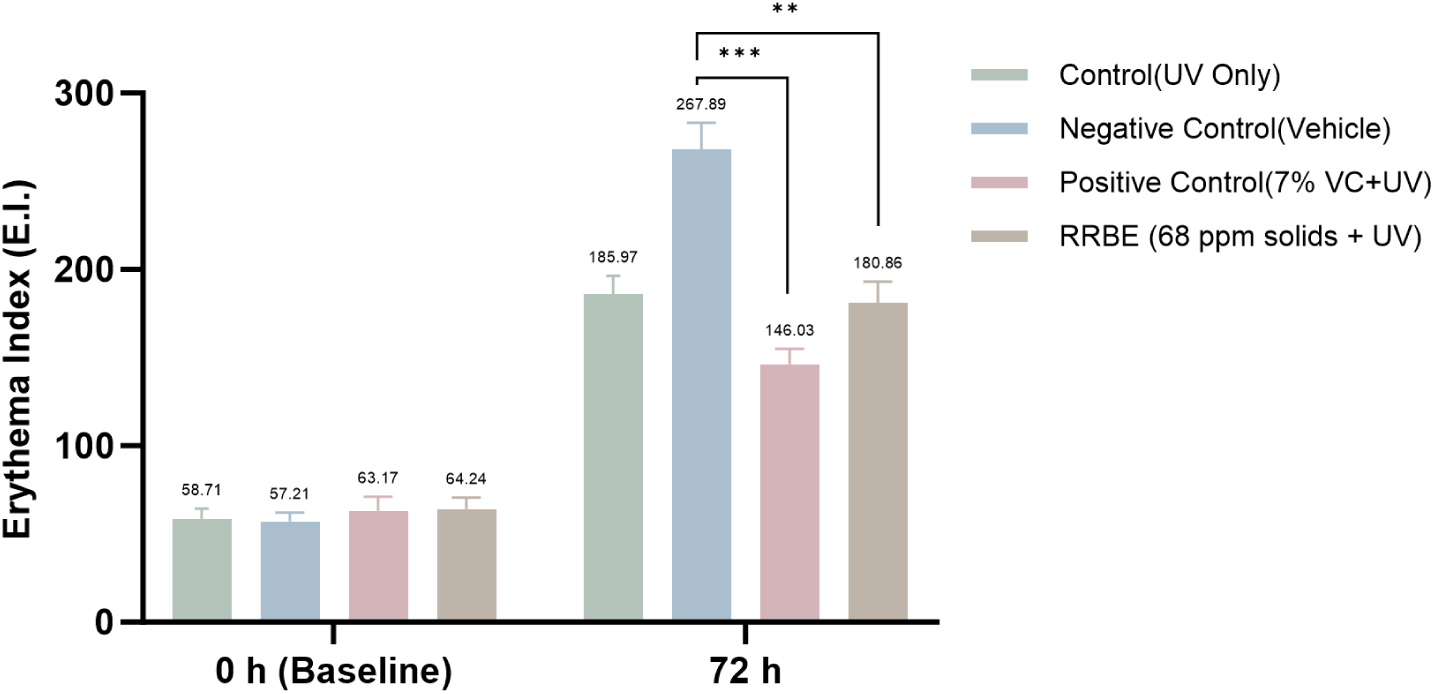
RRBE accelerates the resolution of UV-induced erythema at 72 h post-irradiation. The graph shows the quantitative analysis of the absolute Erythema Index (E.I.) in human subjects (*n* = 10) at 72 h. Control (UV), Negative Control (vehicle + UV), Positive Control (7% Vitamin C + UV), and RRBE (68 ppm solids + UV). Data are Mean ± SD. ^∗∗^*p <* 0.01, ^∗∗∗^*p <* 0.001 vs. Negative Control.

In contrast, treatment with RRBE demonstrated a robust therapeutic effect by significantly accelerating the regression of erythema. At 72 hours, the E.I. in the RRBE-treated group dropped significantly to ∼180.9 (*p <* 0.01 vs. Negative Control), representing a marked recovery from the acute photodamage. Notably, this reduction profile was statistically comparable to that of the Positive Control group treated with 7% Vitamin C (E.I. ∼ 146.0). These data confirm that RRBE possesses potent anti-inflammatory properties *in vivo*, effectively promoting the resolution of UV-induced skin erythema.

#### 3.1.2 RRBE Inhibits UVB-Induced Tissue Damage and Inflammation in 3D Epidermal Models

To substantiate the clinical findings and explore the underlying tissue-level mechanisms, we assessed the protective effects of RRBE in a 3D reconstructed human epidermal model. We first investigated the primary mechanisms of photodamage at the DNA level. We began by assessing the formation of cyclobutane pyrimidine dimers (CPDs), the most critical form of direct DNA damage caused by UVB radiation (Figure 3A). As expected, the Negative Control group (UVB model) showed a massive induction of CPDs, with the mean IOD surging to approximately 32 compared to a negligible baseline of ∼1 in the Unirradiated Control group. Treatment with the Positive Control (Vitamin E, VE) effectively blocked this DNA damage, suppressing the mean IOD to approximately 9. The RRBE-treated group also demonstrated profound photoprotection, significantly inhibiting the formation of these DNA lesions (mean IOD reduced to ∼21), thereby preventing the accumulation of direct DNA damage.

**Figure 3:**
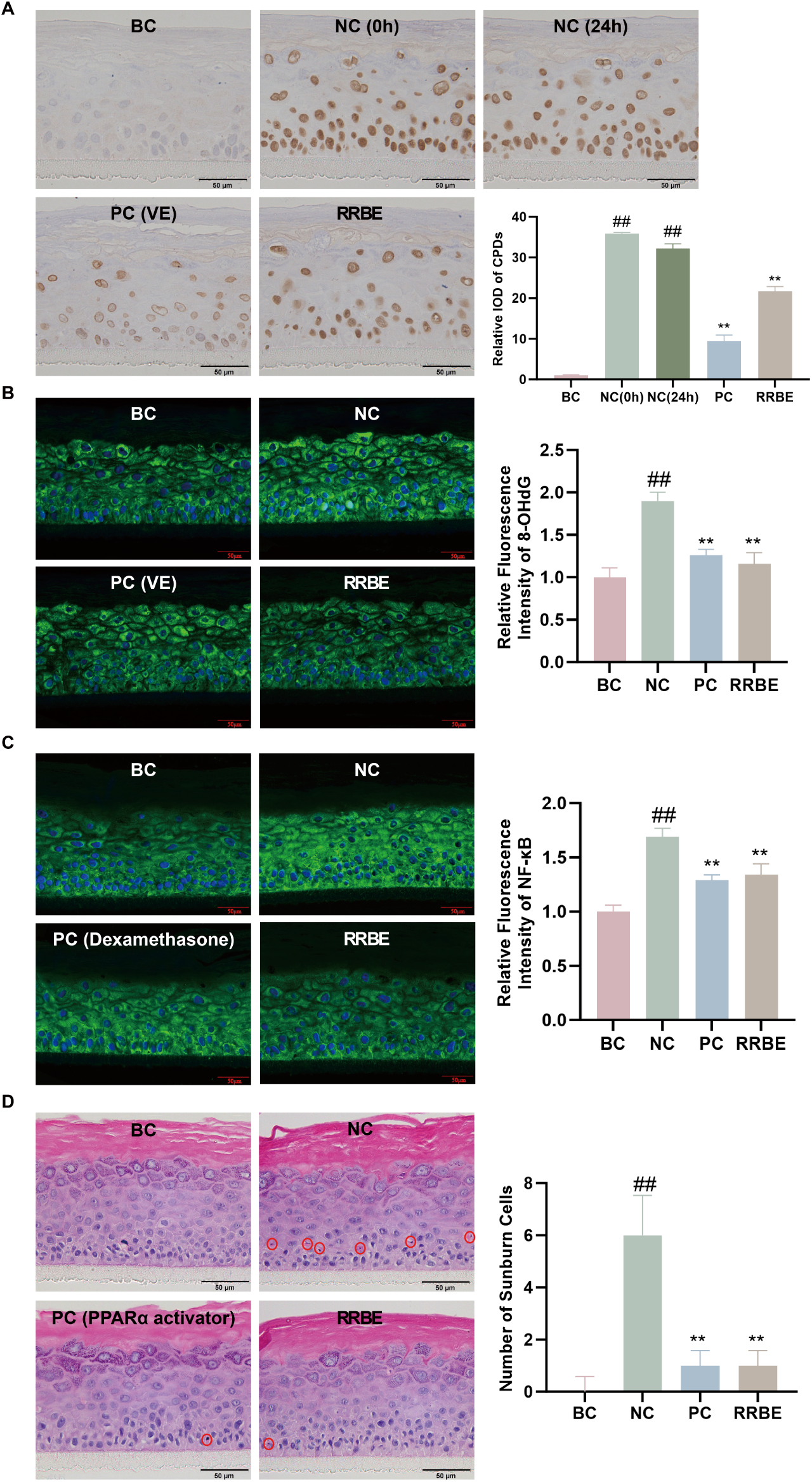
RRBE mitigates UVB-induced photodamage in 3D epidermal models. (**A**) Immunohistochemistry (IHC) for CPDs (Direct DNA damage). RRBE reduces the primary molecular lesions caused by UVB. (**B**) Immunofluorescence (IF) for 8-OHdG (Oxidative DNA damage). RRBE alleviates oxidative stress-induced nucleotide damage. (**C**) IF for NF-*κ*B (Inflammatory signaling). RRBE suppresses the downstream inflammatory transcription factor cascade. (**D**) H&E staining and quantification of Sunburn Cells (SBCs). RRBE ultimately pre-vents acute apoptosis and tissue damage. DAPI (blue) counterstain used for (B, C). Bar graphs represent Mean ± SD (*n* = 3). BC: Blank Control; NC: Negative Control (UVB + vehicle). PC: Positive Control (Vitamin E for A and B; Dexamethasone for C; PPAR*α* activator WY-14643 for D). RRBE: UVB + 1% RRBE. Scale bars = 50 *µ*m. *^##^p <* 0.01 vs. BC; ^∗∗^*p <* 0.01 vs. NC.

Concurrently, we measured the formation of 8-hydroxy-2’-deoxyguanosine (8-OHdG), a primary marker of oxidative DNA damage (Figure 3B). The Negative Control group exhibited a nearly two-fold increase in 8-OHdG levels compared to the Unirradiated Control (Relative IOD rising from 1.0 to ∼1.9). Notably, RRBE treatment effectively reversed this oxidative stress, reducing 8-OHdG levels back to near-basal levels (Relative IOD ∼1.15), demonstrating an antioxidant efficacy comparable to, or slightly exceeding, that of the Positive Control (VE, Relative IOD ∼1.25).

To connect these upstream molecular damage events to the downstream inflammatory response, we evaluated the expression of the pro-inflammatory transcription factor NF-*κ*B (Figure 3C). UVB irradiation caused a strong upregulation of NF-*κ*B expression in the Negative Control group, rising to approximately 1.7 times the baseline level. RRBE treatment markedly attenuated this response, suppressing NF-*κ*B expression to a level (Relative IOD ∼1.35) comparable to that of the potent anti-inflammatory drug Dexamethasone (Positive Control, Relative IOD ∼1.3). This suppression suggests that RRBE effectively blocks the signal transduction pathway linking DNA damage to tissue inflammation.

Finally, we evaluated the ultimate impact on tissue integrity and cell survival by quantifying the formation of Sunburn Cells (SBCs), a hallmark of acute photodamage-induced apoptosis (Figure 3D). Following UVB exposure, the number of SBCs increased dramatically from a negligible baseline (∼0) to approximately 6 cells per field in the Negative Control group, indicating severe acute apoptosis. Strikingly, RRBE treatment effectively rescued the tissue from this lethal injury, reducing the SBC count down to ∼1 cell. This profound reduction highlights RRBE’s potent cytoprotective capacity, achieving an anti-apoptotic efficacy statistically indistinguishable from that of the Positive Control (WY-14643, ∼1 cell).

Taken together, these 3D model data support that RRBE provides comprehensive photo-protection. It mitigates phototoxicity by suppressing key underlying mechanisms—both direct (CPDs) and oxidative (8-OHdG) DNA damage—which subsequently dampens the inflammatory cascade (NF-*κ*B) and ultimately prevents acute cell death (SBCs).

#### 3.1.3 RRBE Mitigates UVA-Induced Oxidative Stress and UVB-Induced DNA Damage Response

To determine whether RRBE confers broad-spectrum protection against photodamage, we first evaluated its efficacy in mitigating oxidative stress in a cellular environment using UVA-irradiated HaCaT keratinocytes. UVA radiation is a primary inducer of reactive oxygen species (ROS), which drive photoaging. As shown in Figure 4A, UVA exposure triggered a sharp accumulation of intracellular ROS (indicated by DCF fluorescence). Pre-treatment with RRBE significantly suppressed this ROS surge, demonstrating its potent capacity to neutralize UVA-mediated oxidative stress at the cellular level.

**Figure 4:**
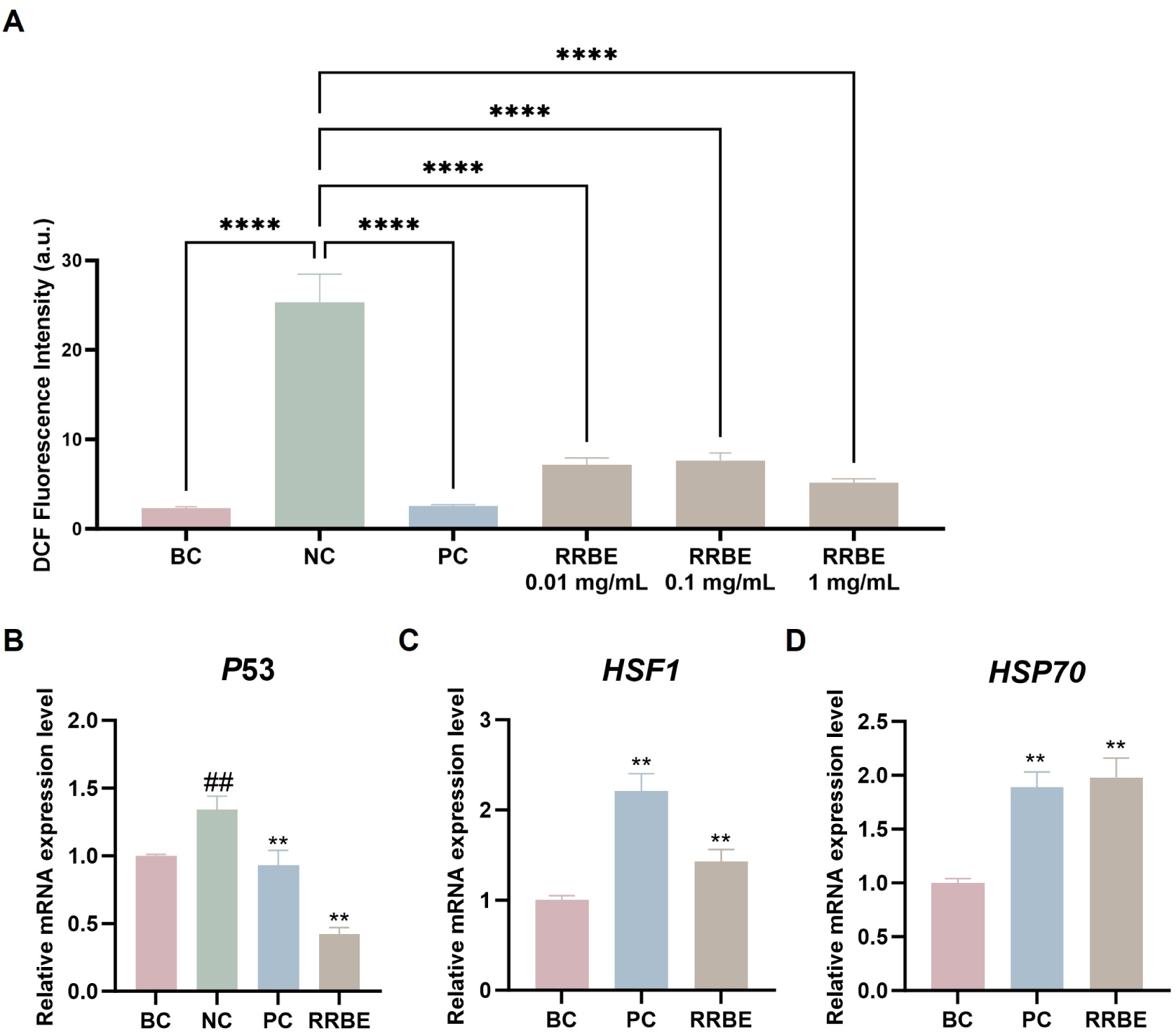
RRBE provides broad-spectrum protection by suppressing oxidative stress and enhancing stress response. (**A**) Intracellular ROS levels in UVA-irradiated HaCaT cells, quantified by DCFH-DA fluorescence. RRBE significantly inhibits UVA-induced ROS accumulation. (**B**) Relative p53 expression (Fold Change) after UVB irradiation. RRBE suppresses the DNA damage response. (**C**) Relative HSF1 expression (Fold Change). (**D**) Relative HSP70 expression (Fold Change). Data are Mean ± SD (*n* = 3). BC: Blank Control; NC: Negative Control (Model); PC: Positive Control; RRBE: RRBE-treated. *^##^p <* 0.01 vs. BC; ^∗∗^*p <* 0.01 vs. NC.

Beyond oxidative stress, we investigated RRBE’s capacity to mitigate UVB-induced genotoxicity. We measured the activation of p53, a critical sensor of the DNA damage response triggered by severe lesions (Figure 4B). Following UVB irradiation, the Negative Control group exhibited a sharp activation of p53, with expression surging to approximately 1.34-fold compared to the non-irradiated control (*p <* 0.01). In contrast, pre-treatment with RRBE markedly attenuated this response, significantly suppressing p53 expression to 0.42-fold (*p <* 0.001), representing a downregulation rate of 68.66%. This brings p53 levels closer to the baseline, confirming that RRBE effectively mitigates direct DNA damage.

Furthermore, we examined the cellular stress response pathway, which is critical for managing protein damage. As shown in Figure 4C, RRBE treatment caused a substantial upregulation of HSF1 (Heat Shock Factor 1) expression, increasing it by 43.00% (to 1.43-fold, *p <* 0.01). Consistent with the role of HSF1 as a transcriptional regulator, RRBE also significantly increased the mRNA levels of its downstream target HSP70 to ∼1.98-fold (*p <* 0.01), mirroring the effect of the positive control (Figure 4D). This upregulation of the HSF1/HSP70 axis demonstrates that RRBE acts as an active modulator, enhancing the intrinsic protein repair machinery.

Finally, we confirmed that the concentrations of RRBE used throughout all cellular assays were non-cytotoxic, as determined by cytotoxicity assays and morphological observation (Supplementary Fig. S1).

### 3.2 Integrated Phytochemical Profiling and Network Pharmacology Reveals Bioactive Modules

To elucidate the material basis of RRBE and systematically map its biological mechanisms, we employed an integrated workflow combining high-resolution phytochemical profiling with deep learning-based target prediction and functional annotation.

We began with a comprehensive composition analysis using LC-MS/MS, which identified a total of 210 compounds (listed in Supplementary Table S2). To systematically organize these components based on their chemical structural features, we constructed a Compound Similarity (CS) network (Tanimoto coefficient ≥ 0.5). Applying the Louvain algorithm partitioned this network into 10 distinct structural modules. As detailed in Table 2, these modules categorize the complex extract into distinct chemical families, primarily encompassing flavonoids, phenolic acids, fatty acids, and nucleosides.

**Table 2:**
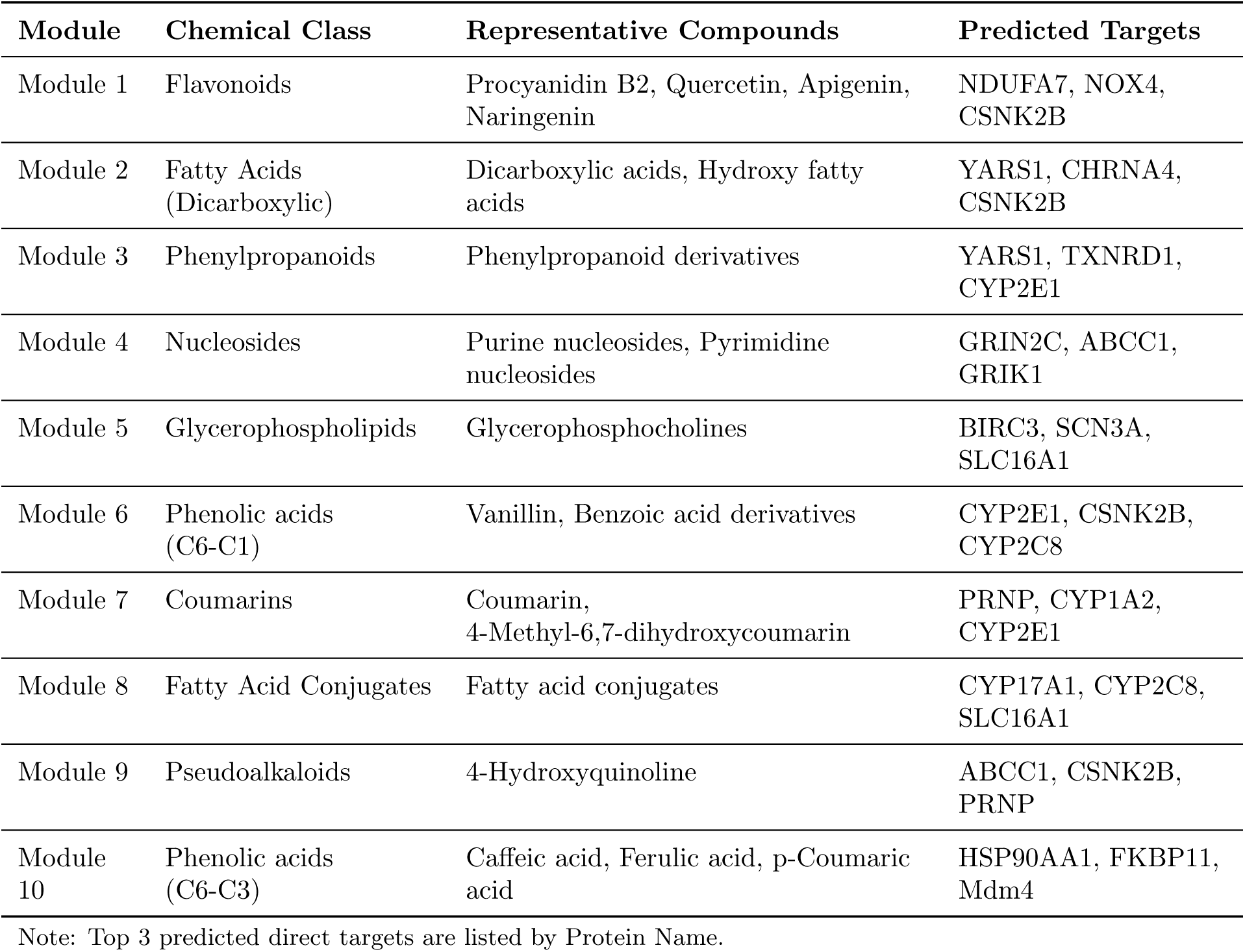
Chemical Composition and Key Predicted Targets of RRBE Bioactive Modules.

To systematically map the specific biological roles of these chemical clusters, we performed global target prediction across all 10 modules using SCOPE-DTI. The prediction analysis revealed that Module 1 and Module 10 emerged as the most pharmacologically significant clusters relevant to photoprotection, exhibiting distinct target profiles (Table 2).

Specifically, Module 1, structurally identified as a flavonoid-rich cluster, was predicted to target key oxidative stress regulators, including NDUFA7, NOX4, and CSNK2B. In contrast, Module 10, which is highly enriched with phenolic acids, showed a strong affinity for proteins involved in stress responses and protein homeostasis, particularly HSP90AA1, FKBP11, and Mdm4. This functional divergence suggests a synergistic mechanism where the flavonoid module targets mitochondrial oxidative stress, while the phenolic acid module manages cellular stress responses.

To statistically quantify the biological relevance of these predicted targets, we performed functional annotation using MetaCore’s pathway database. We specifically analyzed six key dimensions related to photoaging, ordered as follows: DNA damage, oxidation, inflammatory factors, DNA repair, melanin synthesis, and collagen metabolism. The regulatory strength of each module was quantified using a two-tier scoring system that integrated protein importance and pathway prevalence. As illustrated in the radar chart (Figure 5A), distinct functional profiles emerged that were correlated with the direct targets identified above. Consistent with its targets (e.g., NDUFA7, NOX4), Module 1 exhibited exceptional specificity for oxidation and inflammatory factor pathways. Conversely, Module 10, driven by targets such as HSP90AA1, demonstrated a focused convergence on protein homeostasis and stress response pathways. This comparison highlights a functional dichotomy where flavonoids primarily tackle oxidative damage, while phenolic acids manage cellular stress responses.

**Figure 5:**
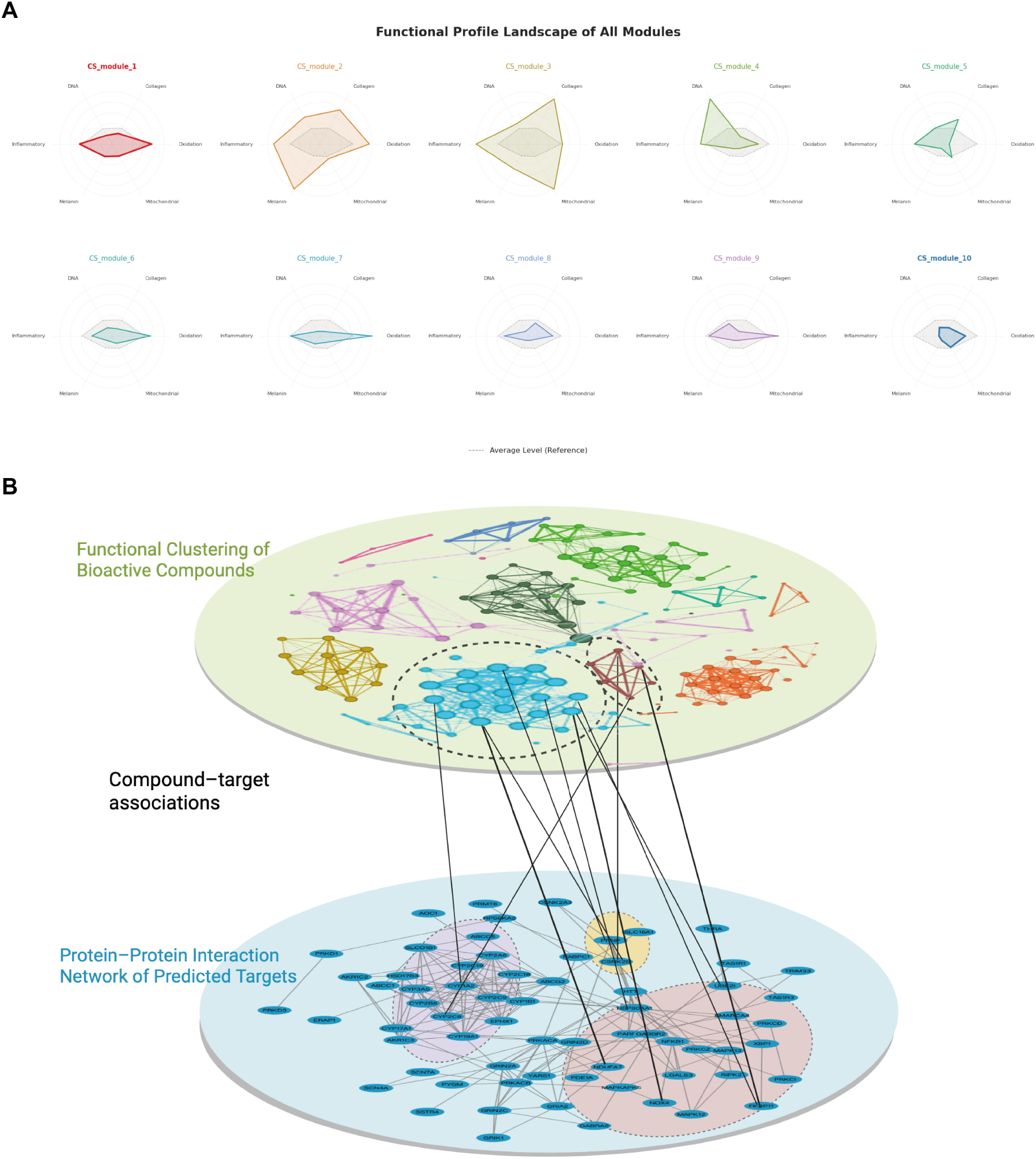
Integrative functional and network analysis of RRBE. (**A**) Module Functional Analysis Comparison (Radar Chart). A radar chart illustrating the pathway importance scores for each RRBE module. Module 1 (flavonoids) shows high specificity for oxidation/inflammation, while Module 10 (phenolic acids) targets stress responses. (**B**) Integrative dual-layer network view linking functional clusters of bioactive compounds (top) to the protein–protein interaction network of their predicted targets (bottom). Colored communities in the upper layer represent compound modules based on their chemical structure. Vertical edges indicate compound–target associations predicted by SCOPE-DTI.

Synthesizing these functional insights with the chemical clustering, we constructed a dual-layer integrative network (Figure 5B) to visualize the polypharmacological landscape of RRBE. The upper layer displays the 10 chemical modules, colored by their chemical structure, while the vertical edges represent the predicted small molecule-protein interactions connecting compounds to the lower layer—the network of protein targets. This global view confirms that RRBE’s efficacy is driven by specific “structure-function” modules: flavonoids (Module 1) regulating mitochondrial oxidative stress, and phenolic acids (Module 10) managing thermal and protein folding stress.

### 3.3 Structural Basis of Target Engagement: *In Silico* Molecular Docking

To provide a structural rationale for the compound-target associations predicted by our network analysis, we performed *in silico* molecular docking to visualize the binding feasibility of the key ligands prior to experimental validation (Figure 6).

**Figure 6:**
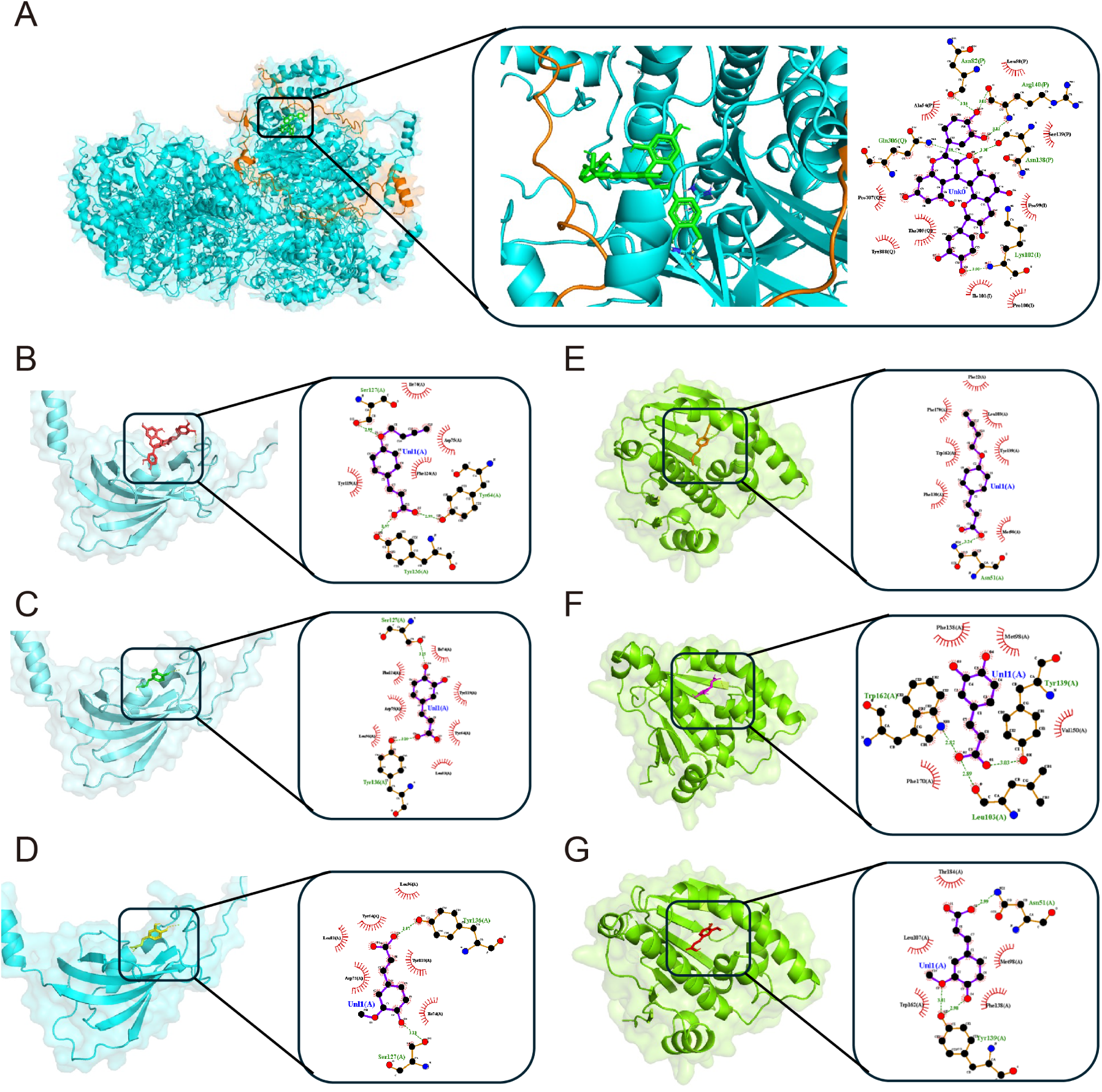
*In Silico* Structural Validation of Key Compound-Target Interactions. (**A**) Predicted binding pose of Procyanidin B2 (Module 1) with the *NDUFA7* subunit of Mitochondrial Complex I (Score: −9.0 kcal/mol). (**B**-**D**) Binding poses of Module 10 phenolic acids with *FKBP11* : (**B**) p-Hydroxycinnamic acid, (**C**) Caffeic acid, and (**D**) Ferulic acid, showing a con-served binding pocket. (**E**-**G**) Binding poses of the same three phenolic acids with *HSP90AA1* : (**E**) p-Hydroxycinnamic acid, (**F**) Ferulic acid, and (**G**) Caffeic acid. Solid lines indicate hydrogen bonds; dashed lines indicate hydrophobic interactions.

We first investigated NDUFA7, a key subunit of the mitochondrial NADH: ubiquinone oxidoreductase (Complex I), which catalyzes electron transfer to drive ATP synthesis. We docked procyanidin B2 (Module 1) against the entire Complex I, defining the docking box to enclose the NDUFA7 subunit (marked in orange). The simulation predicted a high-affinity interaction with a strong binding score of −9.0 kcal/mol (Figure 6A). LigPlot analysis indicated this stability is derived, in part, from a hydrogen bond with residue Lysine-102 of the NDUFA7 subunit, suggesting RRBE may regulate Complex I activity by structurally engaging NDUFA7.

Next, we docked the core phenolic acids from Module 10 against their targets. For FKBP11, a PPIase protein family often targeted by immunosuppressive drugs, docking referenced the known FK506 binding pocket. p-Hydroxycinnamic acid (p-Coumaric acid) showed the highest affinity (binding score: −5.1 kcal/mol), followed closely by ferulic acid (−5.0 kcal/mol) and caffeic acid (−4.9 kcal/mol) (Figure 6B-D). The narrow energy range (ΔΔ*G* = 0.2 kcal/mol) suggests these compounds occupy a similar binding pocket, consistent with the known structural plasticity of the FKBP family.

In contrast, the binding of these acids to HSP90AA1 showed greater variation: p-Hydroxycinnamic acid demonstrated the strongest binding force (−8.0 kcal/mol), significantly outweighing ferulic acid (−6.9 kcal/mol) and caffeic acid (−6.8 kcal/mol) (Figure 6E-G). These favorable binding energies provide a strong structural basis for the plausibility of these direct interactions, supporting our computational predictions and paving the way for biophysical verification.

### 3.4 Biophysical Validation of Key Compound-Target Interactions via CETSA-MS

Following the computational confirmation of binding feasibility, we rigorously verified the direct physical interactions in a physiological environment using CETSA-MS. We prioritized the top-ranked candidates: procyanidin B2 (representing Module 1) and three core phenolic acids (caffeic acid, ferulic acid, and p-coumaric acid, representing Module 10). Assays were conducted using a mixed-cell lysate system comprising keratinocytes (HaCaT), fibroblasts (HSF), and melanocytes (MNT-1) to ensure biological robustness.

The CETSA-MS results provided definitive experimental evidence for the “compound-target” axes. For Module 1, procyanidin B2 induced a substantial thermal stabilization of NDUFA7 (Figure 7A), exhibiting a significant thermal shift (Δ*T_m_* = 12.5^◦^C) with high fitting correlations (*R^2^* = 0.85*/*0.67). As NDUFA7 is a critical subunit of Mitochondrial Complex I, this physical binding confirms the molecular mechanism for Module 1’s role in mitigating mitochondrial oxidative stress.

**Figure 7:**
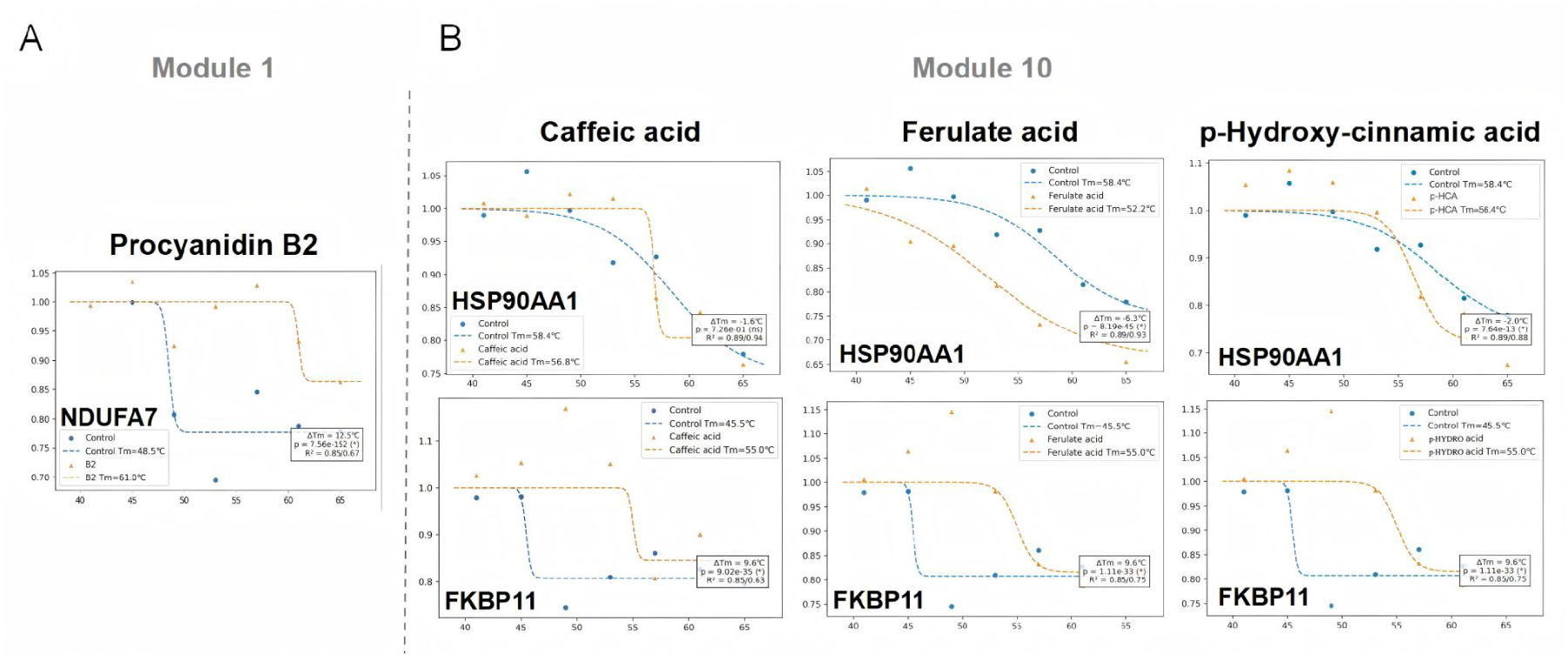
Biophysical validation of target engagement using CETSA-MS. (**A**) Representative thermal melt curves for Module 1: Procyanidin B2 significantly stabilizes *NDUFA7*, confirming mitochondrial target engagement. (**B**) Representative curves for Module 10: Caffeic acid, Ferulic acid (labeled as Ferulate acid), and p-Coumaric acid (labeled as p-Hydroxy-cinnamic acid) all exhibit profound thermal engagement with *HSP90AA1* and *FKBP11*. Note that while FKBP11 shows stabilization, HSP90AA1 exhibits modulatory thermal shifts, validating them as conserved stress-response targets. Data points represent normalized protein abundance at indicated temperatures (*n* = 3).

Concurrently, Module 10 validation revealed distinct engagement profiles for the core phenolic acids (Figure 7B). Caffeic acid demonstrated a robust stabilization of FKBP11 (Δ*T_m_* = 9.6^◦^C, *R^2^* = 0.85*/*0.63) while displaying a modulatory thermal shift on HSP90AA1 (Δ*T_m_*= −1.6^◦^C, *R^2^* = 0.89*/*0.94). Similarly, ferulic acid exhibited strong binding to FKBP11 (Δ*T_m_* = 9.6^◦^C, *R^2^* = 0.85*/*0.75) and a destabilizing shift on HSP90AA1 (Δ*T_m_* = −6.3^◦^C, *R^2^* = 0.89*/*0.93). Furthermore, p-coumaric acid showed a consistent targeting pattern, stabilizing FKBP11 (Δ*T_m_* = 9.6^◦^C, *R^2^* = 0.85*/*0.79) and modulating HSP90AA1 (Δ*T_m_*= −2.0^◦^C, *R^2^* = 0.89*/*0.88). These biophysical data not only corroborate the accuracy of our SCOPE-DTI predictions and docking simulations but also experimentally confirm that RRBE’s bioactivity is driven by the synergistic modulation of distinct mitochondrial (NDUFA7) and stress-response (FKBP11/HSP90AA1) targets.

## 4 Discussion

Ultraviolet (UV) radiation is a major extrinsic driver of skin aging and inflammation. It generates damage through several mechanisms, including direct DNA lesions, oxidative stress, and dysregulated immune responses [27, 28]. Although natural products rich in polyphenols have long been used for photoprotection, their inherent complexity makes it difficult to move from empirical observations to a clear mechanistic understanding. In this study, we established an integrated, AI-driven framework combining phytochemical profiling, network-based target prediction (SCOPE-DTI), and target validation using CETSA-MS. This approach enabled systematic dissection of the multi-target mechanisms of RRBE, a natural complex derived from red rice bran.

At the phenotypic level, RRBE showed strong photoprotective activity in both clinical and experimental models. In human subjects, RRBE significantly reduced UV-induced erythema, indicating effective suppression of acute vasodilation and inflammation. In the 3D reconstructed epidermal model, RRBE reduced key markers of direct and oxidative DNA damage, including cyclobutane pyrimidine dimers (CPDs) and 8-hydroxy-2’-deoxyguanosine (8-OHdG). It also reduced the formation of sunburn cells and clearly suppressed NF-*κ*B activation. These findings suggest that RRBE does more than act as a simple antioxidant. It is capable of blocking major UV-induced damage pathways at the tissue level.

Further mechanistic insights were obtained from 2D HaCaT keratinocyte assays. RRBE markedly reduced UVB-induced p53 activation, suggesting that it helps alleviate severe stress r-lated to DNA damage. Additionally, RRBE robustly activates the HSF1/HSP70 pathway, which supports enhanced protein homeostasis and increases cellular resilience in response to stress. These findings align with recent research highlighting the broad antioxidant, anti-inflammatory, and pigmentation-modulating potential of pigmented rice bran extracts in mitigating UV-induced skin damage [29].

Through LC-MS/MS profiling and compound similarity networking, we deconstructed RRBE into distinct structural clusters, among which Module 1 (flavonoids) and Module 10 (phenolic acids) were highlighted as the primary bioactive drivers. While core compounds within these modules, such as proanthocyanidins and caffeic acid, have been previously reported to exhibit significant photoprotective properties [30, 31, 32], their specific molecular targets had remained elusive. To bridge this gap, we combined AI-driven prediction (SCOPE-DTI), biophysical validation (CETSA-MS), and structural simulation (molecular docking) to systematically map their targets. First, we identified the accessory subunit NDUFA7 of mitochondrial Complex I as a key target of the flavonoid module. Although classified as a non-catalytic accessory protein, NDUFA7 is involved in maintaining Complex I stability and regulating ROS generation [33]. Recent functional studies have further shown that the loss of NDUFA7 reduces Complex I activity and impairs mitochondrial function [34]. Decreased Complex I activity leads to increased ROS production and further mitochondrial dysfunction [35]. UV exposure can damage Complex I, increase mitochondrial ROS, and subsequently activate NF-*κ*B–mediated inflammatory signaling [36, 37]. Our data show that procyanidin B2 stabilizes NDUFA7, which reduces ROS generation at its source and explains why RRBE strongly suppresses ROS and NF-*κ*B activity. Earlier literature often reported that flavonoids improve mitochondrial oxidative balance [38, 39]. However, most studies did not specify the exact protein target. Our findings provide target-level resolution by identifying NDUFA7 as a direct and functionally relevant site of action.

In parallel, we discovered a second pathway at the endoplasmic reticulum involving FKBP11. FKBP11 is an ER-resident chaperone that regulates the unfolded protein response (UPR). Recent work demonstrates that FKBP11 modulates the PERK-NRF2 axis, enabling cells to adopt more adaptive responses under chronic ER stress [40]. Our results indicate that phenolic acids in RRBE bind directly to FKBP11 and may enhance its chaperone function. This activity can help relieve protein-folding stress caused by UV and ROS. The effect also complements the mitochondrial pathway. While RRBE reduces ROS production at the mitochondrial level, it also enhances the ER’s ability to mitigate the consequences of oxidative stress.

A third mechanism involves HSP90AA1, a central regulator of inflammatory and antioxidant signaling. HSP90 participates in multiple pathways, including NF-*κ*B activation, NLRP3 inflammasome regulation, and NRF2-mediated antioxidant responses [41, 42]. Our findings show that phenolic acids in RRBE bind strongly to HSP90AA1, and CETSA-MS confirmed enhanced thermal stability. It is known that HSP90 maintains the stability of the IKK complex, which is required for NF-*κ*B activation. Pharmacological inhibition of HSP90 disrupts IKK stability, prevents I*κ*B degradation, and blocks NF-*κ*B nuclear translocation [35]. This mechanism is consistent with the NF-*κ*B suppression observed in the 3D model.

Taken together, our results support a multi-layer regulatory network through which RRBE provides protection. At the mitochondrial level, RRBE limits ROS production by stabilizing NDUFA7. At the ER level, RRBE enhances protein-folding homeostasis through FKBP11. At the inflammatory level, RRBE regulates NF-*κ*B and NLRP3 through its interaction with HSP90AA1 (Figure 8). This integrated model closely aligns with the network pathology of photoaging, which involves oxidative stress, DNA damage, proteostasis imbalance, and chronic inflammation. It also helps explain why complex natural mixtures often show stronger *in vivo* protection than single antioxidants.

**Figure 8:**
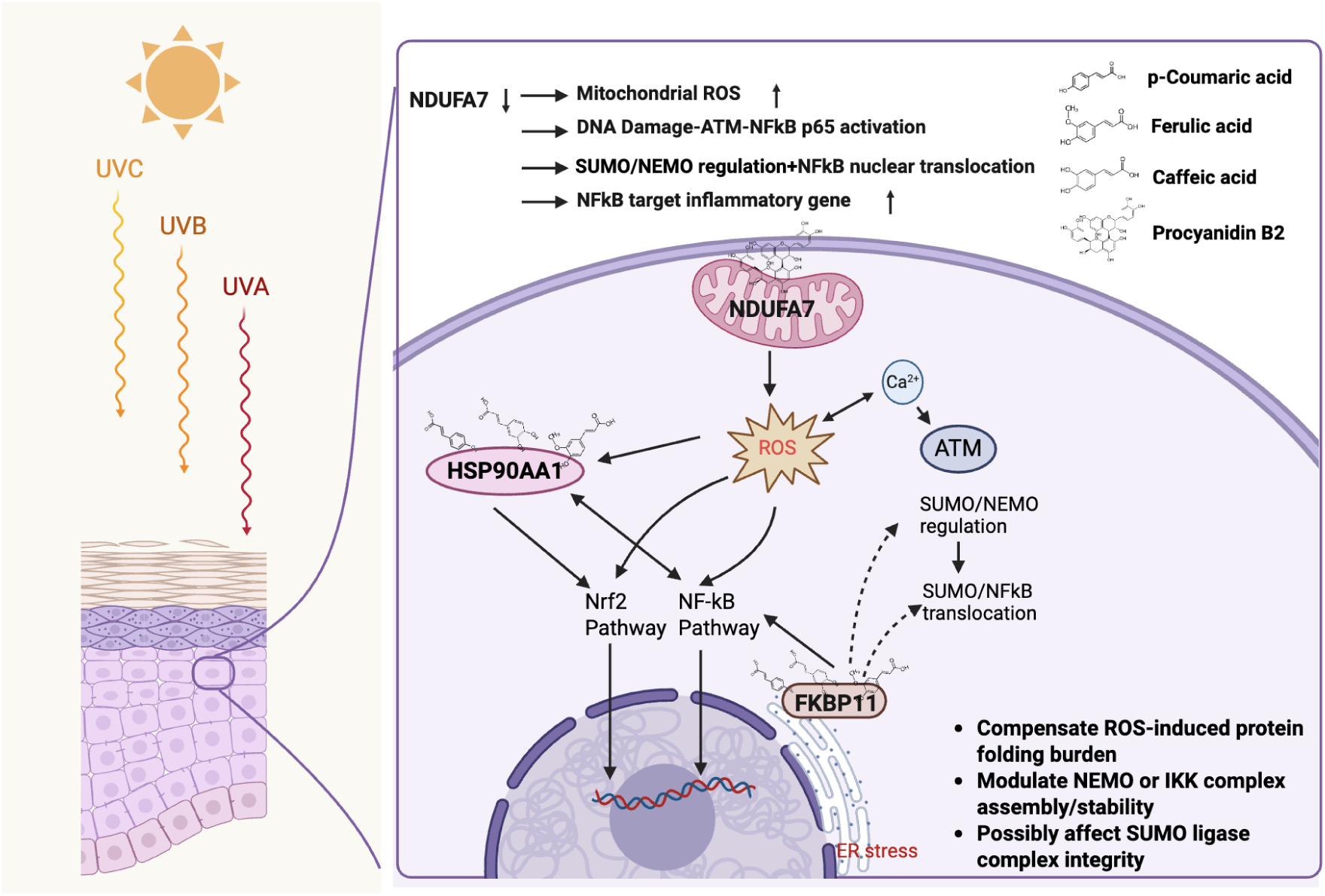
Mechanistic Hypothesis Construction. RRBE employs a multi-target strategy: Module 1 targets NDUFA7 to prevent mitochondrial ROS bursts, while Module 10 tar-gets FKBP11 and HSP90AA1 to resolve ER stress and modulate inflammatory signaling (NF-*κ*B/Nrf2), providing comprehensive protection against photoaging.

From a methodological perspective, this study demonstrates the advantage of combining deep learning–based target prediction with CETSA-MS validation. CETSA-MS, however, has inherent limitations related to detection sensitivity and restricted temperature windows. As a result, RRBE’s full target spectrum may not yet be completely captured. Additional experiments, including chemical proteomics and CRISPR-based genetic perturbation, will be necessary to further validate the complete target network. Another limitation is that we used representative anchor compounds for each module and did not systematically examine whether components within the same module act synergistically or antagonistically. Future work involving multi-component combinations, module-level CETSA-MS, and functional genetic studies will help clarify intra-module interactions and strengthen causal evidence for the mechanisms.

Despite these limitations, the findings of this study have important implications for photo-protection research and the development of natural products. Overall, our data support the characterization of RRBE not merely as a simple antioxidant, but as a bioactive agent capable of engaging and coordinating multiple cellular stress-response systems. Furthermore, the integrated platform established here provides a reproducible and systematic roadmap for advancing natural product research. This approach facilitates the transition of botanical complexes from traditional empirical use toward mechanism-guided, evidence-based development.

## 5 Conclusion

In conclusion, this study proposes the polypharmacological mechanisms underlying the photo-protective efficacy of RRBE, a natural complex derived from red rice bran. Through an integrated workflow combining LC-MS/MS profiling, AI-based target prediction, CETSA-MS and molecular docking validation, we identify flavonoids and phenolic acids as the major functional modules driving RRBE’s activity. These components act through three key and complementary stress-response pathways: stabilization of NDUFA7 to suppress mitochondrial ROS generation, engagement of FKBP11 to alleviate ER protein-folding stress, and modulation of HSP90AA1 to maintain inflammatory homeostasis. Together, these coordinated actions provide a systems-level protective framework against UV-induced damage. Beyond mechanistic insight into RRBE, this work establishes a scalable “AI-guided, cell-validated” discovery pipeline that can be broadly applied to study complex natural products and guide the rational design of next-generation, mechanism-driven photoprotective and anti-aging formulations.

## 6 Author Contributions

T.S.L., Y.G.C., X.L., T.Z., Y.C.D.L., and H.D.H. conceived the study. Data collection was conducted by T.S.L., X.L., X.J., S.H.H., J.Y.L., L.P.L., H.L.Z., S.F.L., J.L., H.X.Z., X.T.Z., Y.F.W., S.Q.N., Z.H.Z., and Y.G. T.S.L., Y.G.C., and X.L. designed the methodology. Computational analysis was performed by Y.G.C. and H.Y.H. Wetlab experiments were executed by T.S.L., X.L., X.J., and S.H.H. The original manuscript was drafted by T.S.L., Y.G.C., X.L., Y.C.D.L., and H.D.H. All authors reviewed and approved the final manuscript.

## 7 Funding

This work was financially supported by the Shenzhen Science and Technology Program (JCYJ2022-0530143615035, JCYJ20250604141235046, JCYJ20250604141041017); the National Natural Science Foundation of China (No. 32070674, No.82272568); the Warshel Institute for Computational Biology funding from Shenzhen City and Longgang District (LGKCSDPT2025001); Shenzhen-Hong Kong Cooperation Zone for Technology and Innovation (HZQB-KCZYB-2020056, P2-2022-HDH-001-A); Guangdong Young Scholar Development Fund of Shenzhen Ganghong Group Co., Ltd. (2021E0005, 2022E0035); Phase III Government Matching Fund of Shenzhen Ganghong Group Co., Ltd. (2023E0012); Guangdong Science and Technology Program (2024A0505050001, 2024A0505050002); 2023 The Second Affiliated Hospital of the Chinese University of Hong Kong, Shenzhen Joint Fund Project (HUUF-MS-202308, HUUF-MS-202309); CUHK(SZ) HOMEY HEALTH Microbiome and EndoMetabolic Digital Health Research Center (2024E0049); CUHK(SZ) GeneBioHealth Advanced Molecular Diagnostics Laboratory (2024E0088); and the Better Way Group – Chinese University of Hong Kong (Shenzhen) Warshel Joint Laboratory for skin health and active molecule innovation (2024E0087); The Guangdong Basic and Applied Basic Research Foundation (2022A1515220168).

## 8 Data Availability Statement

Data are contained within the article.

## 9 Acknowledgments

This work was supported by the Warshel Institute for Computational Biology funding from Shenzhen City and Longgang District (LGKCSDPT2025001). The Institute is recognized as the Guangdong Provincial Key Laboratory of Digital Biology and Drug Development. We would like to express our sincere gratitude to the Vincent & Lily Woo Foundation for their generous support of the Vincent & Lily Woo Fellowship in Memory of Albert Wong. This fellowship, endowed by the Vincent & Lily Woo Foundation, is provided through MCMIA Foundation Limited, and we are deeply grateful for their invaluable contribution to our research. We also acknowledge the collaborative support from Shaanxi BioCell General Testing Co., Ltd. We specifically thank Yimeng Wang and Yuhan Jiang for conducting the human efficacy evaluations, Fanghui Sun for performing the cellular assays, and Qi Sun for preparing the formulation samples. Finally, we wish to express our appreciation to Yue Liu for the essential support in project management and timeline coordination, which greatly facilitated the completion of this work.

## 10 Conflicts of Interest

T.S.L., Y.G.C., X.L., S.Q.N., Z.H.Z., Y.G., T.Z., Y.C.D.L., and H.D.H. are listed as inventors on a patent application related to the work presented in this manuscript (Patent Application No. 202512056758.8). The remaining authors declare no competing interests.

## Notes

### Summary of Updates

This revision has been submitted to update the Declaration of Interests section. The authors have amended the statement to disclose a patent application (Application No. 202512056758.8) related to the work presented in this manuscript. Specifically, authors T.S.L., Y.G.C., X.L., S.Q.N., Z.H.Z., Y.G., T.Z., Y.C.D.L., and H.D.H. are listed as inventors. No changes have been made to the scientific data, figures, or conclusions.

